# Relationship between Brain and Body Temperature in Anesthetized Animals Measured by Ultralocal Thermometry

**DOI:** 10.1101/2024.12.08.627392

**Authors:** A.A. Osypov, V.K. Krokhaleva, A.M. Romshin, I. Yu Popova

## Abstract

Temperature is one of the least studied biophysical characteristics of the brain, although both the rate of biochemical reactions and the electrical activity of nervous tissue directly depend on it. The purpose of this study was to analyze the relationship between brain and body temperatures in anesthetized animals. Simultaneous measurement of brain and body temperature (rectally) was carried out at ambient temperature controlled with a thermal mat. The temperature of the deep layers of the somatosensory cortex was measured by ultralocal thermometry using a diamond thermometer. Under anesthesia, the body and brain temperature dropped to 27 C (4 degrees above ambient temperature). When the thermal mat was turned on, the brain and body began to heat up synchronously. The brain initially lagged behind, but when the critical temperature was reached, it began to release heat in quantities exceeding the influx from the blood, reaching physiological values of 37C. A reverse experiment with decreasing the temperature of the thermal mat showed a similar picture: a synchronous start of the decrease, but with reverse dynamics. Thus, we can distinguish two phases of the brain’s reaction to external heating: passive - when neuronal activity is decreased, and active - after internal regulatory mechanisms are triggered, which, in its turn, slows down the temperature drop. In general, the data obtained in the present work indicate that the temperature of neural tissue is not linearly related to body temperature.

## Introduction

Brain temperature is a fundamental but virtually unexplored biophysical parameter. The brain shows exceptional sensitivity to disturbances in temperature balance, rapidly enhancing ischemic or other neurological damage (Minamisawa et al., 1990), and a prolonged increase in body temperature by as little as 1 degree significantly impairs cognitive abilities (Piil et al., 2020). The relationship between changes in brain temperature and various parameters of neuronal activity, blood-brain barrier permeability, glial cell activity, brain ionic and water balances, and the structural integrity of various brain cell types has been shown (Moser et al., 1993). The brain is also known to be highly metabolically active, and all energy used for brain metabolism is ultimately converted into heat (Kiyatkin, 2019). However, the matter of the brain temperature as a biophysical parameter reflecting neural activity and affecting various neural functions remains to be unknown. This is primarily due to the fact that until recently there were no direct and accurate methods for measuring brain temperature. Almost all brain temperature data have been obtained either by indirect estimation (Kiyatkin, 2019) or by using methods with low spatial and temperature resolution (Marshall et al., 2006). However, it is clear that accurate, high-resolution temperature characterization of brain tissue can provide unique information about the fundamental mechanisms underlying brain function, helping to define new ways to diagnose pathological conditions and treat brain diseases.

In the last decade, significant progress has been made in the development of miniaturized temperature-sensitive elements, which has significantly advanced biological thermometry at the subcellular level (Ishiwata et al., 2014; Yang et al., 2014; Okabe et al., 2012; Romshin et al., 2022). However, as for in vivo experiments on model animals, there are only unique pairs of works on direct local thermometry of nervous tissue in the literature (Fedotov et al., 2020; Romshin et al., 2024; Sui et al., 2023), leaving this topic practically unexplored.

The present study analyzed the dynamics of the somatosensory cortex temperature and rectal body temperature of BALB/c mice simultaneously under anesthesia under controlled ambient temperature, both at room temperature and when the body was heated with a heating mat.

## MATERIALS AND METHODS

### Experimental animals

Experiments were performed on BALB/c mice (25-33 g) obtained from the Stolbovaya Laboratory Animal Nursery (https://www.pitst.ru/, Moscow region, Chekhov district, Stolbovaya settlement). Animals were kept in cages with unrestricted access to food and water, in a temperature-controlled room (22°C ± 1°C) and a 12-hour light/dark cycle. The protocol was approved by the Committee on Bioethics of Animal Experiments of the Institute of Theoretical and Experimental Biophysics of the Russian Academy of Sciences.

Experiments were performed on anesthetized animals (n=9). General anesthesia was created by intraperitoneal injection of urethane (650mg/kg) and xylazine (50mg/kg). The head was fixed in stereotaxis (Kopf Instruments), the skull surface was thoroughly cleaned of skin and connective tissue, and a hole was drilled in the skull to insert the thermometer. The exposed brain surface was protected from desiccation with a drop of medical glycerol.

### Experimental design

In anesthetized animals, simultaneous recording of rectal temperature and temperature in the somatosensory cortex of the brain was performed at room temperature (23-24°C) and when the animal was heated to physiological level (37°C) using an animal heating mat (14W power, NomoyPet, China).

### Registration of the brain temperature

Deep somatosensory cortex temperature was measured by ultrolocal thermometry by diamond heat thermometer provided by Wonder Technologies (https://wondercvd.com).

The thermometer comprises a glass microcapillary in whose inner channel a single 20 μm diamond microparticle is integrated and an optical fiber is introduced to excite and read luminescence from the microdiamond (Romshin et al., 2021). The fabricated thermometer was fixed on a high-precision 3D micromanipulator and inserted into the primary somatosensory cortex to a depth of 0.8-1.1 mm at coordinates AP =-0.5-1.0; L=-1.5-2.0 (Paxinos and Franklin, 2001). The approximate area of temperature acquisition was about 100 μm^3^.

### Registration of the body temperature

A standard veterinary digital electronic thermometer TA-288 (Russia) was used to measure body temperature *per rectum*. Recordings were made in 10 seconds increments.

## Results and discussion

### Brain and rectal temperature in mice under anesthesia

Under xylazine-urethane anesthesia, the rectal body temperature of mice fell to 27.2±1 °C, which is 10 degrees below the physiological norm, and insignificantly depends on the room temperature, being 3±1 °C above it. The somatosensory cortex temperature at the surgery zone was 26.3±1.2 °C, insignificantly 1±1 °C lower than the body temperature.

Temperature measurements were started after complete immersion of animals in general anesthesia. Rectal temperature of experimental animals was stable at 27.2±1 °C, which is in agreement with previously published data describing the phenomenon of anesthesia-induced hypothermia for different types of anesthesia and for different animal species (Lenhardt, 2010; Murphy & Murnane, 2019). The dependency on the room temperature, though insignificant, shows the balance between the lowered physiological heat production and the ambient heat exchange.

Brain somatosensory cortex temperature at the surgery zone was 26.3±1.2 °C, which is only insignificantly 1±1 °C lower than the body temperature, indicating the cooling effect of the scalping and the surgery hole in the skull. Thus, in the state of anesthesia, the brain temperature decreases by at least 11 °C compared with the temperature recorded in the brains of mice in the state of active wakefulness

Our data on rectal temperature are in good agreement with the literature sources, while fundamentally complementing them due to the registration of ultralocal brain temperature in experimental animals. As far as we concern, such a huge temperature fall in the brain has been never reported for the *in vivo* experiments.

### Dynamics of changes of the brain and rectal temperature in anesthetized mice during body heating with a thermal mat

Turning on the thermal mat on which the animal was lying caused a simultaneous rise in rectal temperature and brain temperature. Interestingly, the dynamics of the body and brain temperatures increase differed greatly (see fig. 1 and 2).

**Fig. 1.**
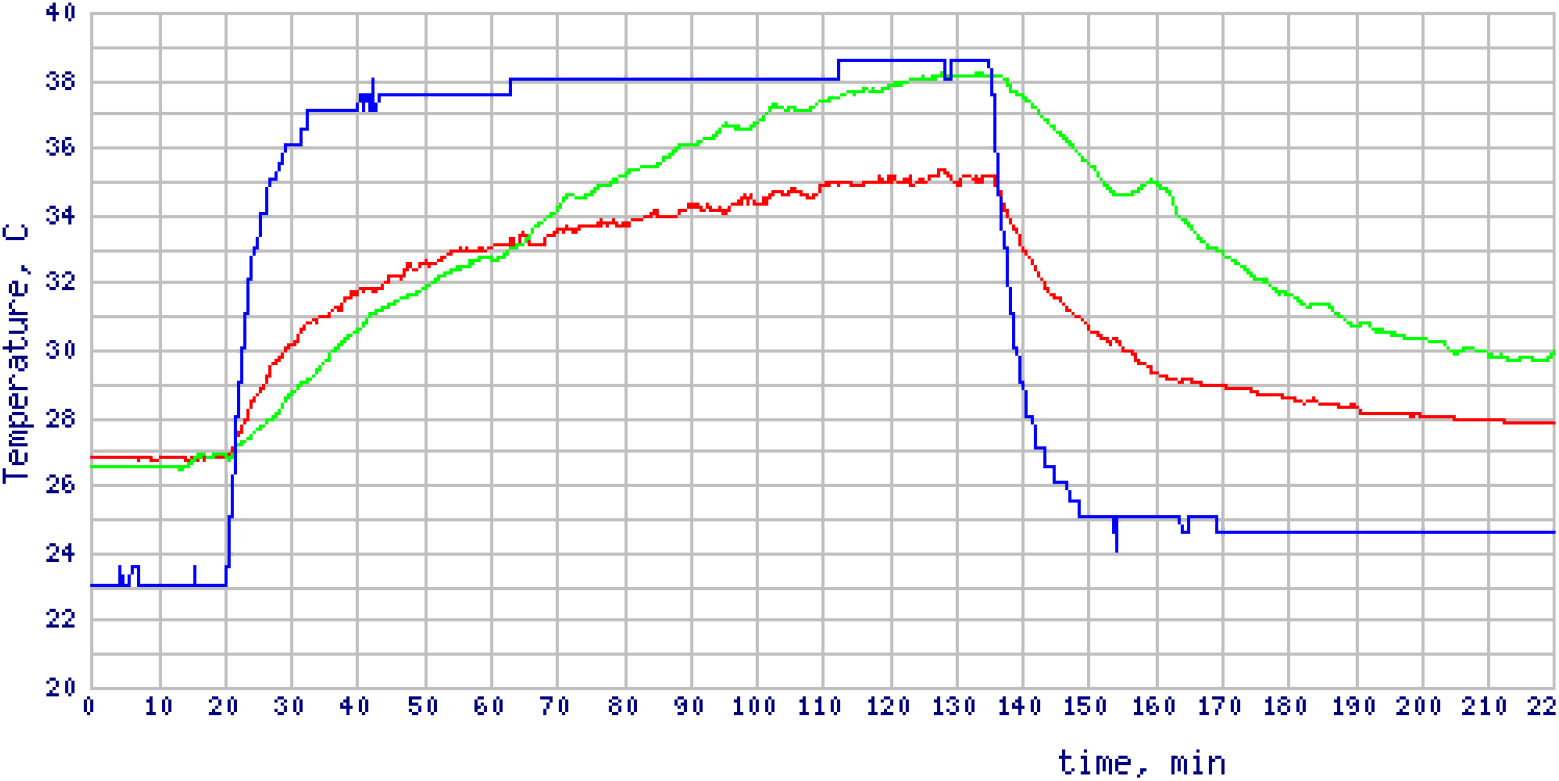
Body temperature (rectally), brain temperature and heating mat temperature. The mat was turned on at 20 minutes of the experiment and turned off at 135. On the right edge, from top to bottom: brain (green), body (red) and mat (blue) temperatures.

**Fig. 2.**
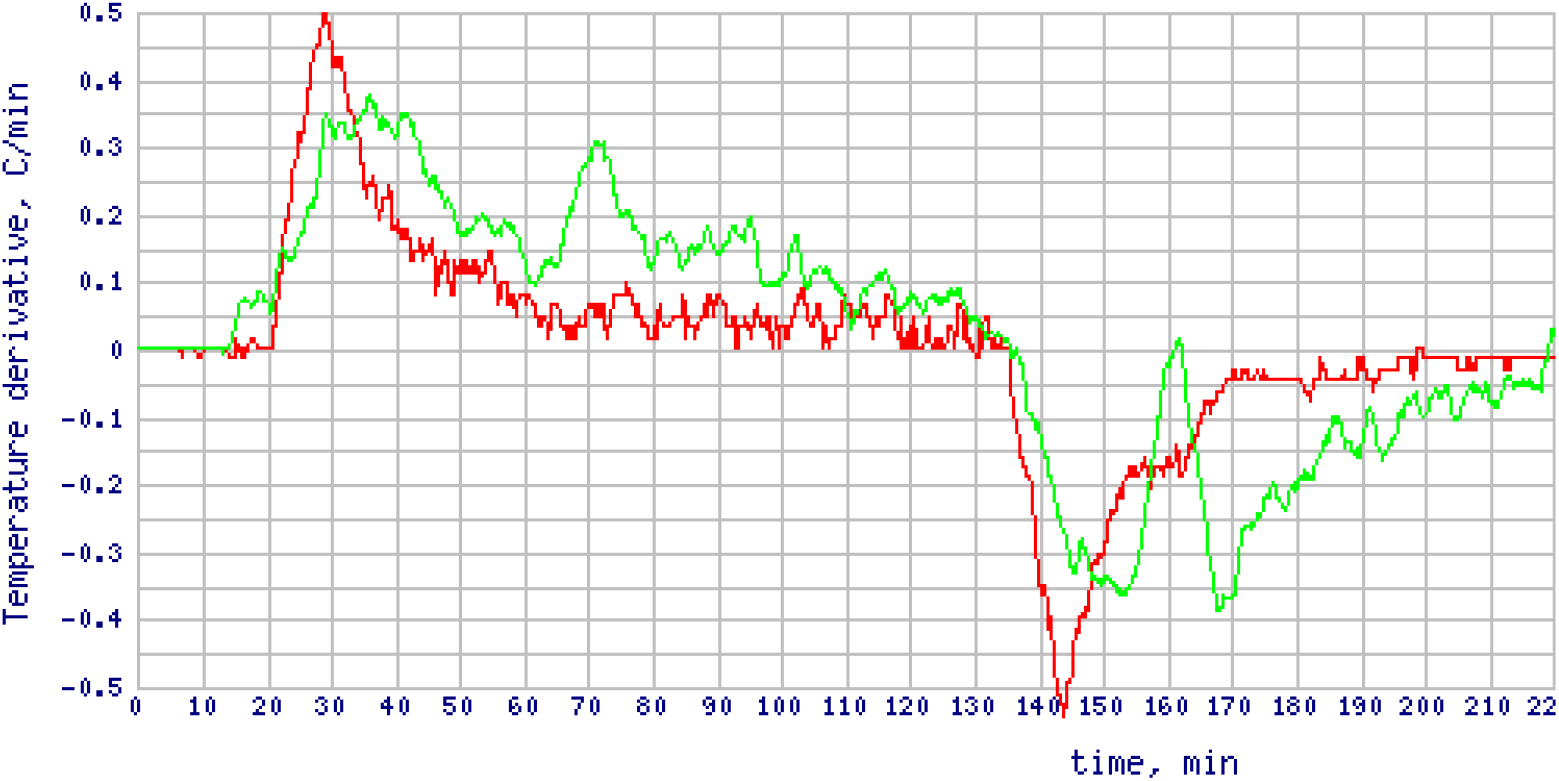
Dynamics of the body and brain temperature changes, expressed as the temperature change upon time in degrees Celsius per minute. Brain temperature, green, bottom line at 30 min; body rectal temperature, red, top line at 30 min.

Twenty minutes after turning on the heating mat, rectal temperature and brain temperature increased by 4.6±1.7 and 3.6±1.4 °C, respectively.

The body began to heat up rapidly, but gradually the rate of temperature increase slowed down and reached a plateau during prolonged heating. The rectal temperature never rose to the mat temperature of 36.3±1.6 °C on which the animal was lying. On average, the maximum of the rectal temperature was 33±1 °C at 3 degrees below that of the thermal mat.

Brain temperature also began to rise immediately after turning on the heating mat, but at a slower rate than body temperature, and at 20 minute increased only by 3.6±1.4 °C. Thus, the difference in brain and rectal temperatures increased by a degree, so at the initial stage of heating the temperature rise in the brain lags behind that of the body.

However, further on the brain temperature began to rise more rapidly than the body one and at 38±12 minutes they reached the cross at 32.4±0.8 °C. Here from the pure data variation we see that the brain temperature rather than the heating time obviously was the leading parameter. When the heating prolonged further, the brain heated up by itself up to 38.1 °C, that is 3 degrees above the rectal temperature and belongs to the normal physiological realm. The average difference was 1.7±1.3 °C.

The moment of the brain internal heating turns on may be figured out by the cross between the curves of the temperatures growth rate that occurs at 29.3±0.6 °C at 12.2±0.8 minute. Obviously, at that point the temperature rise rate in the brain equals that of the body and might be still attributed to the passive blood heat distribution. However, as the initial heating rate in the brain lagged behind that of the body, we can assume that the active phase of the brain heat production occurs even earlier. Actually, the brain internal heat production turns on somewhere near the inflection point, but we have not enough resolving power in our data to confidently estimate its position.

Thus, we can distinguish two phases of the brain’s reaction to external heating: the passive one under the reduced neuronal activity and the active - after the internal regulatory mechanisms are triggered on. The passive phase seems to be maintained by the heat provided by the blood warmed by the heating mat, while the active phase indicates the triggering of the brain internal heating through signaling mechanisms, rather than as a result of blood flow effects. We can only speculate on these mechanisms being of the neurons activation of some kind, but the exact underlying physiological and biochemical machinery remain to be discovered and explored.

### Dynamics of changes of the brain and rectal temperature in anesthetized mice after termination of heating with a thermal mat

The reverse experiment with the mat turning off showed similar dynamics of temperature decrease - synchronized onset of body and brain temperature decrease, but with different dynamics. Both when the temperature rises and falls, the body responds faster, but then the brain takes over.

The rectal temperature passively followed the heat distribution by the mat, and when it is switched off, rapidly drops initially with the gradual decline of the decrease rate with a plateau at the end, directly mirroring the heating dynamics, thus providing the idea of the overall passive body temperature response. The brain temperature however, already starting from higher levels, drops slower and more linear and stays higher for quite some time, indicating prolonged heat-producing activated state.

Interestingly, in one mouse the brain temperature continued to grow even after 15 minutes the mat was turned off, reaching 35.5 °C at 43 minutes, that is 3 degrees more than the rectal maximum of 32.5 °C at the switch-off time and striking 5.5 °C more than the synchronous rectal temperature. Obviously this indicates a kind of a self-sustaining active physiological process of the yet unknown nature.

## Conclusion

Currently, fiber optic thermometers allow high-resolution dynamic temperature measurements in the living brain compared to even the most accurate implantable thermistors and thermocouples [22, 23]. Using this new methodological approach in this pilot study, simultaneous dynamic measurements of local temperature in the somatosensory cortex of the brain and rectal body temperature of anesthetized mice were performed under the controlled ambient temperature conditions.

Under xylazine-urethane anesthesia, the rectal body temperature of mice fell to 27 °C that is 10 degrees below the physiological norm, and insignificantly depends on the room temperature, being 3 degrees above it. The somatosensory cortex temperature at the surgery zone was insignificantly 1 degree lower than the body temperature, indicating the cooling effect of the scalping and the surgery hole in the skull. Thus, in the state of anesthesia, the brain temperature decreases by at least 11 °C compared with the temperature recorded in the brains of mice in the state of active wakefulness. As far as we concern, such a huge temperature fall has been never reported for the *in vivo* brain.

After the external heating with a mat began, both temperatures started to rise with the body temperature leading at the first phase so that the difference in brain and rectal temperatures increased by a degree after 20 minutes of heating. Gradually the rate of the body temperature increase slowed down and reached a plateau during prolonged heating. The rectal temperature never rose to the mat temperature nor came close to the physiological norm, being only 33 °C on average.

However, from the 12th minute the brain temperature began to rise more rapidly than the body one and somewhere at 25-40 minutes they reached the cross at 32 °C, with the brain temperature rather than the heating time obviously being the leading parameter. When the heating prolonged further, the brain heated up by itself up to 38.1 °C, that is 3 degrees above the rectal temperature and belongs to the normal physiological realm.

Interestingly, the difference between the brain and body temperatures varies greatly and can reach as much as 3 and even 5.5 degrees celsius. The physiological causes and consequences of such behavior may be of great importance to the brain functioning in norm and pathology.

The reverse dynamics occurs after the heating mat turning off - the body temperature rapidly drops initially with the gradual decline of the decrease rate with a plateau at the end, directly mirroring the heating dynamics, thus providing the idea of the overall passive body temperature response. The brain temperature however, already starting from higher levels, drops slower and more linear and stays higher for quite some time, indicating prolonged heat-producing activated state.

Thus, we can distinguish two phases of the brain’s reaction to external heating: the passive one under the reduced neuronal activity and the active - after the internal regulatory mechanisms are triggered on. The passive phase seems to be maintained by the heat provided by the blood warmed by the heating mat, while the active phase indicates the triggering of the brain internal heating through signaling mechanisms, rather than as a result of blood flow effects. We can only speculate on these mechanisms being of the neurons activation of some kind, but the exact underlying physiological and biochemical machinery remain to be discovered and explored.

In general, the data obtained in the present work indicate that the temperature of neural tissue is not linearly related to body temperature. Studies of temperature characteristics of nervous tissue in different functional states can make a significant contribution to the understanding of the fundamental mechanisms underlying brain functions in norm and pathology.

## Acknowledgments

We thank the Company Wonder Technologies (https://wondercvd.com) for providing the diamond heat thermometry system.

## Funding

This work was supported by the Russian Science Foundation (grant №23-14-00129) and by the State Assignment of the Russian Federation № 075-01025-23-01.

## Notes

### Competing Interest Statement

The authors have declared no competing interest.

